# Predicting vertical ground reaction forces from 3D accelerometry using reservoir computers leads to accurate gait event detection

**DOI:** 10.1101/2022.02.14.480318

**Authors:** Margit M. Bach, Nadia Dominici, Andreas Daffertshofer

## Abstract

Accelerometers are low-cost measurement devices that can readily be used outside the lab. However, determining isolated gait events from accelerometer signals, especially foot-off events during running, is an open problem. We outline a two-step approach where machine learning serves to predict vertical ground reaction forces from accelerometer signals, followed by force-based event detection.

We collected shank accelerometer signals and ground reaction forces from 21 adults during comfortable walking and running on an instrumented treadmill. We trained one common reservoir computer using segmented data using both walking and running data. Despite being trained on just a small number of strides, this reservoir computer predicted vertical ground reaction forces in continuous gait with high quality. The subsequent foot contact and foot off event detection proved highly accurate when compared to the golden standard based on co-registered ground reaction forces. Our proof-of-concept illustrates the capacity of combining accelerometry with machine learning for detecting isolated gait events irrespective of mode of locomotion.

## 1 Introduction

Estimating the presence of a step using mobile devices can be realized with fair accuracy and relative ease (1-5). Yet, many details of the stepping cycle remain opaque such as foot contact and foot off moments, but also more detailed gait characteristics, such as loading responses in (ambulant) clinical contexts. In most of the current literature on wearables, event estimations are rule-based and often require searching for an area of interest (6, 7). This is true for data from inertial measurement units but also for data derived from only accelerometers. Algorithms are optimized for either walking (8-15) or running (16-23) and vary depending on sensor location and type, and on speed. As it stands, they may not generalize to other settings.

Machine learning approaches may provide welcome alternatives. They have been employed to predict stepping moments and gait phases by extracting different features recorded from inertial measurement units (24-37), 3D marker data (38-41), electromyography (42), pressure sensors (43), and textile sensors (44). Across the board, though, these approaches required a priori feature extractions and are, hence, potentially jeopardized by selection bias.

Stepping instants can readily be identified using (the shape of) ground reaction forces (GRFs), typically obtained from force plate signals (45, 46). With these, one can specify single/double support and flight phases and, correspondingly, the mode of locomotion, i.e., walking or running. As such it seems obvious to first seek to estimate the GRF’s shape from wearable sensors and to subsequently using these predicted waveforms to determine gait events. Also here, machine learning has been successful. GRFs during the stance phase, for instance, has been estimated using only acceleration (47-50), a combination of acceleration and angular velocity (51-54), or marker-based kinematics (55, 56). The GRFs during double stance phase could be estimated via marker-based kinematics (57, 58) and the GRFs during the full gait cycle using accelerometers placed on the torso (59). Yet, these approaches often appeared tailored to the data under study rendering their generalizability questionable, but more importantly, in almost all cases, they only managed to predict GRFs for the stance phase, whereas the swing phase is of great importance when investigating running.

A recent review revealed that the shape of the GRF can most accurately be estimated from accelerometry (60) and another found neural networks as a promising tool to do so (61). This triggered the idea of estimating vertical GRF waveforms from acceleration signals of the lower extremities via reservoir computers, more specifically via echo state networks (ESNs) (62-64). ESNs are ‘minimal’ forms of recurrent neural networks. Thanks to the reservoir’s ‘complex’ structure, they may come with great computational capacities (65, 66). In the absence of feedback, one can train them with a very simple and robust rule: optimizing output weights by mere linear regression. This is particularly appealing when considering that typical data sets on gait are fairly limited in size and that any implementation of machine learning in wearable devices should come with low computational costs.

In the following, we conceptually prove that a single reservoir computer can accurately predict vertical GRF waveforms from shank accelerometer signals, which allows for detecting gait events during walking and running with particularly high precision.

## 2 Methods

### 2.1 Participants

We included data of 21 healthy young adults (13 male / 8 female) in the analysis with a mean ± standard deviation age, height, and weight of 20.8 ± 1.0 years, 181.7 ± 10.3 cm, 71.1 ± 9.8 kg, respectively. The recorded speeds were 1.24 ± 0.12 m/s for walking and 2.20 ± 0.14 m/s for running. The participants provided written informed consent in compliance with the Declaration of Helsinki. The experimental design was approved by The Scientific and Ethical Review Board of the Faculty of Behavioural & Movement Sciences, Vrije Universiteit Amsterdam, Netherlands (File number: VCWE-2022-008R1).

### 2.2 Experimental protocol

Participants walked and ran at their preferred speeds on an instrumented dual-belt treadmill (Motek Medical BV, Culemborg, Netherlands) in tied-belt mode wearing their own shoes. The preferred walking and running speeds were determined for each participant followed by a familiarization. The preferred speeds were determined by starting at either 2 km/h or 6.5 km/h for walking and running, respectively, and slowly increasing the speed by 0.1 km/h until the participant felt it was comfortable (67). Subsequently, the same process was repeated at a speed 1.5 km/h above this by now slowly decreasing the speed by 0.1 km/h until a new or the same preferred speed was reached. If the two speeds differed more than 0.4 km/h from each other, a third iteration was done, and irrespective of two or three iterations, the mean of the determined preferred speeds was used. The participants were instructed to step with each foot on a separate belt to be able to record the time series of the ground reaction forces from one leg. For each participant a walking and a running trial were recorded of each five minutes in length. Only consecutive strides absent of artefacts (stepping on the wrong belt) were retained leaving an average of 72 strides per trial for analysis (range: 49-116 strides).

Tri-axial accelerometers, built into the probe of the wireless bipolar surface electromyography system (Mini wave plus, Zerowire; Cometa, Bareggio, Italy), were mounted on the right and left tibia, respectively (see Figure 1). The accelerometers were not placed in the same exact position due to inexperienced researchers. Accelerometer data were sampled at 2000/14 Hz with sensitivity of ± 16g. The vertical GRFs were sampled at 1 kHz and re-sampled to 2000/14 Hz.

**Figure 1:**
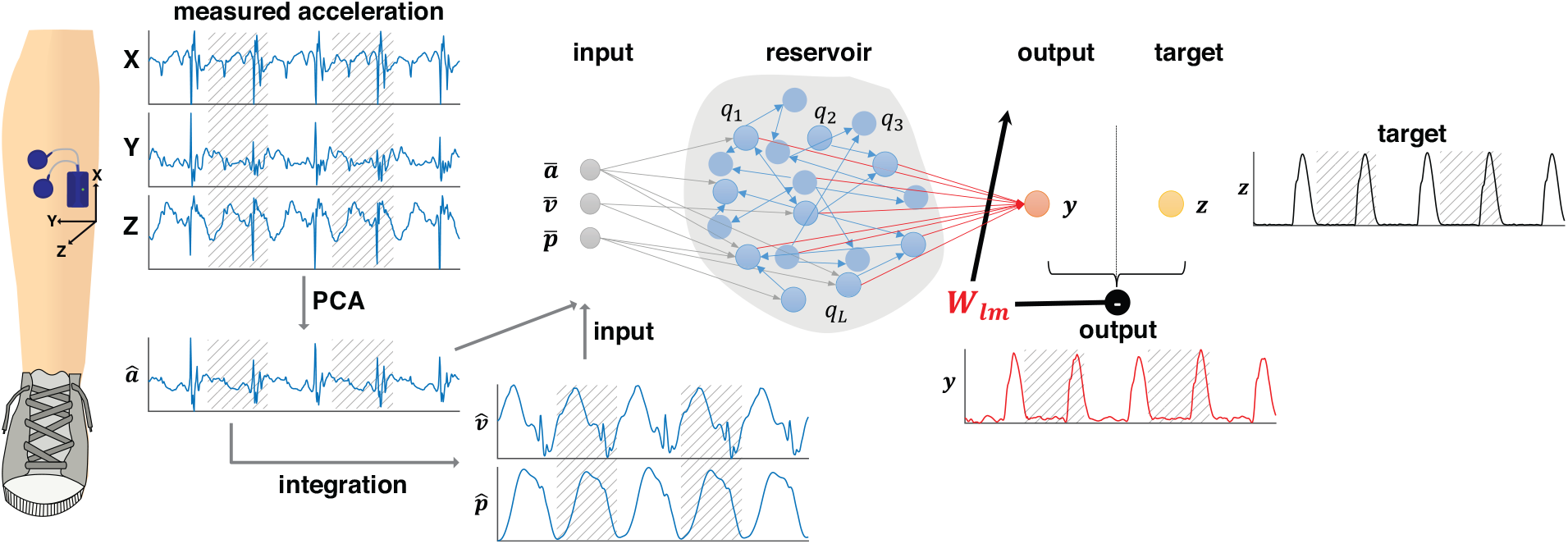
A reservoir computer was implemented to predict the vertical ground reaction forces. Tri-axial accelerometer data were recorded from the shank. The accelerometer data were re-oriented using a principal component analysis (PCA). The first prinicipal component (â) was integrated once to obtain the velocity 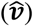 and position 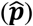 data. The input ***x*** consisting of the normalised accelerometer (***ā***), velocity 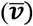, and position 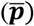 data were subsequently mapped onto the sparsely, randomly connected reservoir ***q***. This reservoir generated the output ***y***, the predicted normalised vertical ground reaction forces (in red), via output weights ***W***. When training, the output was compared to the target ***z***, i.e. the measured vertical ground reaction forces (in black). Minimising the difference between generated output and target served to adjust the weights (denoted here as ***W***_***lm***_). For training, data were segmented into strides, here represented by hatched and unhatched areas. Testing was conducted on continuous data. Data from walking and running were pooled.

A single reservoir computer was trained to predict ipsilateral continuous vertical ground reaction forces based on the shank accelerometer data recorded during walking and running. Figure 1 contains a schematic of the pre-processing steps and the implemented machine learning approach. Further details are presented in the following.

### 2.3 Data processing

Accelerometer signals *a* were first ‘standardized’ to their principal axes using principal component analysis (PCA) (68, 69):

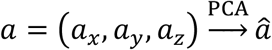

with *â* along the direction of maximum variance and being the only principal component that was retained. *â* was integrated twice over time (after a bi-directional high pass Butterworth filter with cut-off at 1 Hz, 2^nd^ order) to generate likewise standardized velocities 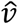 and positional data 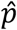 :

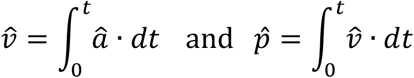

Per subset (trial) these signals were normalized (70) by means of

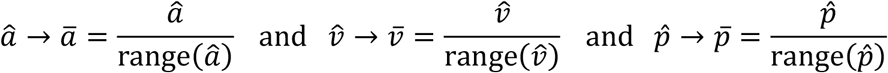

before combining them as three-dimensional input data

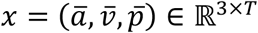

with *T* indicating the number of samples in time. Vertical ground reaction forces *F*_*z*_ were normalized. With this, the target signal for our machine learner (see below) could be defined as z-score (70)

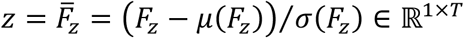

with *μ* and *σ* denoting the mean and standard deviation over time per trial.

Stepping moments (foot contact and foot off events) were identified based on the measured *F*_*z*_ through mere thresholds: First, the *F*_*z*_ was scaled to a range [0 1], then weakly filtered with a polynomial Savitzky-Golay filter (1^st^ order, ±30 ms = in total 9 samples) (71). The foot contact was defined as the last point below a threshold (12.5% of the maximum of the data) nearest the ascend of the *F*_*z*_ of the stance phase; similarly, the foot off was defined as the first point crossing the same threshold nearest the descending *F*_*z*_ (46, 72).

Data were split according to the defined foot off events for further analysis. We considered 36 samples in time on either side of the foot off as transients when correcting for learning errors in the beginning or end of the data. These transients also served to ensure that data were independent of the true events as 36 samples represent different percentage of the stride for walking and running, respectively.

### 2.4 Reservoir computer

We adopted the leaky ESN implementation by Jaeger and Haas (62), (see also 73, 74). In brief, we built a reservoir of *N* nodes *q* = (*q*_1_, *q*_2_, …, *q*_*N*_) ∈ ℝ^*N*×*T*^ that received an input *x* = (*x*_1_, *x*_2_, …, *x*_*k*_) ∈ ℝ^*K*×*T*^ and generated output *y* = (*y*_1_, *y*_2_, …, *y*_*M*_) ∈ ℝ^*M*×*T*^. During the supervised training mode output was compared with target *Z* = (*Z*_1_, *Z*_2_, …, *Z*_*M*_) ∈ ℝ^*M*×*T*^ by means of the L_2_-norm (75); cf. Figure 1.

The reservoir dynamics can be written as

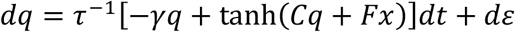

where *C* ∈ ℝ^*N*×*N*^ denotes the connectivity of the reservoir. Here, *C* was set as a sparse, random matrix specified by a sparseness parameter; its weights were normalised for a given spectral radius (the relative magnitude of the leading eigenvalue of *C*). *F* ∈ ℝ^*N*×*T*^ was set to be a dense matrix allowing for an optional scaling of the input values when mapping them onto the reservoir. The quantity *ε* stands for uniformly distributed, uncorrelated noise, i.e., *ε* ∈ *ε*𝒰(−1,1), with *ε* being reasonably small. The output is given by

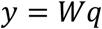

with *W* ∈ ℝ^*M*×*N*^, which is the matrix of the to-be-learned output weights.

Learning was realized by ridge regression, i.e.

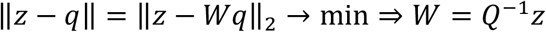

where *Q* = [*q*_1_, …, *q*_*T*_] and (·)^−1^ denotes the pseudo-inverse. In the case of multiple time series, i.e. *S* steps (see below), we defined 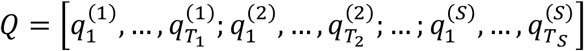 and accordingly we used *Z* = [*Z*^(1)^; … ; *Z*^(*S*)^].

The network parameters were set as follows: *N* = 1000, spectral radiu*s* = 0.5, *F* = [0.1; 0.5], τ = 1, *γ* = 0.5 and *ε* = 10^−4^. The noise was primarily added to minimize the risk of overfitting and we put *ε* = 0 after learning.

#### 2.4.1 Stepping moments from the predicted vertical ground reaction forces

Stepping moment identification of the predicted vertical ground reaction force waveforms was implemented in the same way as for the measured vertical ground reaction forces (see above).

Estimation of gait events such as foot contact and foot off from vertical GRF waveforms are considered the golden standard in movement analysis. An example of the detection algorithm during walking and running can be found in Figure 2. One sample difference between the events based on the measured and predicted vertical GRF waveforms equaled ∼7 ms due to the relatively low sampling frequency, common to wearable accelerometers.

**Figure 2:**
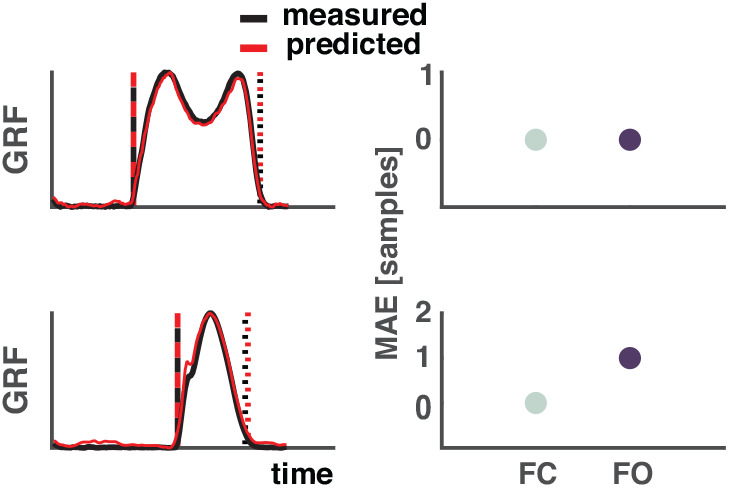
Example of the estimation of foot contact and foot off events from measured and predicted ground reaction forces. Top: walking, bottom: running. Left side: Measured vertical GRF waveforms are depicted in black and the predicted ones in red. The vertical dashed lines represent foot contact (black: measured, red: predicted) and the dotted lines represent foot off (black: measured, red: predicted). Right side: Differences in samples between events based measured and predicted vertical GRF. One sample equal ∼7 ms. Abbreviations: MAE: mean absolute error, FC: Foot contact, FO: Foot off.

### 2.5 Statistical evaluation

The normalized root mean square error *ϵ* and the coefficient of determination *R*^2^ served for quality assessment of the predicted vertical ground reaction forces. We defined them as follows:

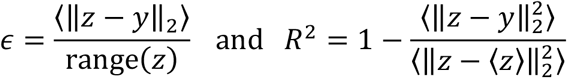

Prediction of stepping moments were validated using the mean absolute error defined as

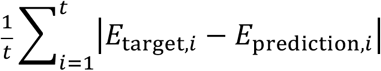

where, *E*_target,*i*_ and *E*_predition,*i*_ refer to target and prediction events *i* = 1, …, *t*, respectively.

We evaluated the training via cross-validation with 50% of the data segmented and subsequently used for training, 25% continuous data for validation, and the remaining 25% continuous data for testing. We performed 100 repetitions with a random draw each time. A continuation rule was used, such that if the *R*^2^ of the validation data were all positive, the testing could be employed, and the training was satisfactory. A maximum of 100 repetitions were allocated for validation and in cases where the validation criteria was not satisfied, the training was stopped. In all cases, the number of strides used from each trial during training was reduced to 25 to ensure a balanced design.

Two scenarios served to assess the robustness of the reservoir computer as sketched in Figure 3.

**Figure 3:**
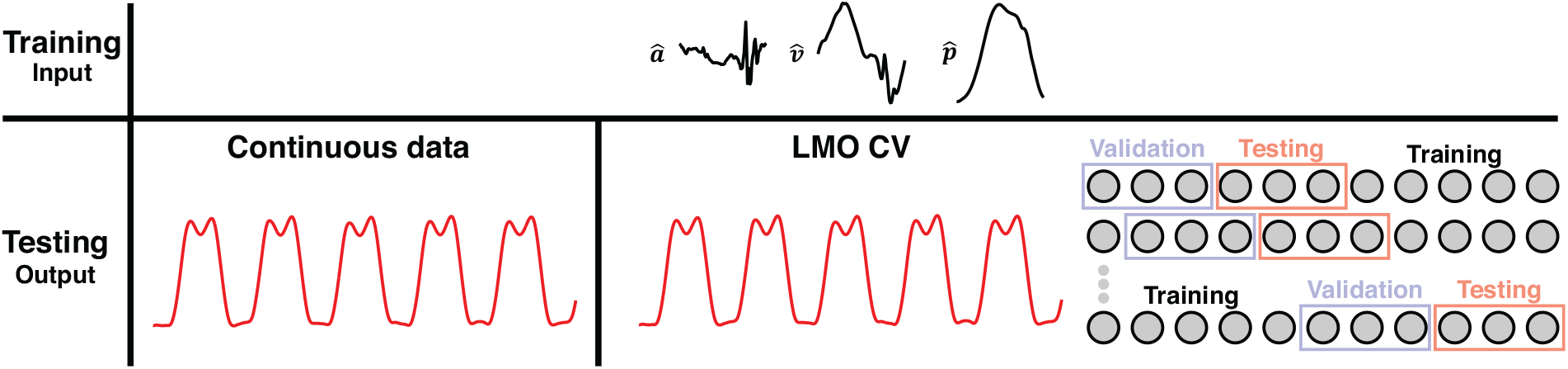
Schematic of the different testing scenarios used for validating the robustness of the reservoir computer. A random trace of a walking trial is shown here. The input data (***â***, 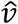, 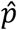) were first segmented into strides (we considered 36 samples in time on either side of the foot off as transients when correcting for any learning errors in the beginning or end of the data). The first scenario, the training is performed on segmented data and the testing on continuous data. The continuous data (in red) represents the output of the reservoir, the vertical GRF waveforms. Secondly, the training was performed on segmented data, testing was done on continuous, but a leave-M-out cross-validation (LMO CV) was employed (split: 50/25/25% for training, validation, and testing, respectively). The vertical GRF in red represents the output of the trained reservoir during testing. Abbreviations: LMO CV: Leave-M-out cross-validation.

#### 2.5.1 Training on segmented data – validating and testing on continuous data

The applicability of our machine learning approach on continuous data were verified by training the reservoir computer on single strides and subsequent testing on continuous data from each trial (see Figure 3). First, we extracted a random 50% of continuous data from each trial before segmenting the remaining data into strides. The continuous data was split in two so 25% of the data were used for validation and testing, respectively. The segmented data were pooled across trials and conditions before being used for training. Training was validated by verifying the mean R^2^ over the validation set to be positive (see above for definition). Whenever validation did not pass with success, training was repeated using the same subset but other randomly chosen initial conditions (here in all cases validation was passed on first attempt). The entire process was repeated 100 times to allow for statistical evaluation as mentioned earlier.

Additionally, we estimated the minimum amount of data needed to secure a good reconstruction quality (*R*^2^ > 0.95), so the training data were reduced. We repeated the training 100 times from 4% of the total dataset to 50% of the total dataset. The validation and test sets remained 25% each for all runs (here the smaller training set sizes required re-learning but eventually validation was passed with success).

#### 2.5.2 Leave-M-out cross-validation

To test the machine learner’s ability to work as a classifier across participants, it was first trained and validated on a subgroup of participants and then tested on others that was unknown to the machine learner. The cross-validation split was performed based on trials. *M* trials were held out and the remaining 42-M trials were split 75/25% of the total dataset for training and validation. A total of 42 repetitions were performed. In the main text we report the result for *M* = 1 while the range *M* = 1,2 …, 6 is depicted in Supplemental Fig. S2.

Unless specified otherwise, means and standard deviations are provided and were calculated as either the grand averages or the standard deviations across the 100 repetitions.

## 3 Results

A total of 3020 strides were included from 42 trials (1249 walking strides [21 trials] and 1771 running strides [21 trials]). Here, we would like to note that we only show results of the right-side analysis, as the left-side results were very similar. Given this similarity one may pool data across sides to increase the sample size but, as will become clear, this was not needed.

The performance of 100 repetitions in predicting GRF waveforms exceeded 95% when combining walking and running data. The coefficients of determination *R*^2^ were 0.96 ± 0.00 and the normalized root-mean squared errors *ϵ* were 6.8 ± 0.3% (mean ± SD); cf. Figure 4. On average, the subsequently extracted foot contact and foot off events deviated from those based on the measured vertical GRF waveforms by 3 and 4 samples. This corresponds to mean absolute errors of 21.9 ± 6.5 ms and 29.1 ± 16.0 ms for foot contact and foot off events, respectively. Here we would like to add that the likewise convincing results when training the network on only walking or on only running are provided as Supplemental Fig. S1.

**Figure 4:**
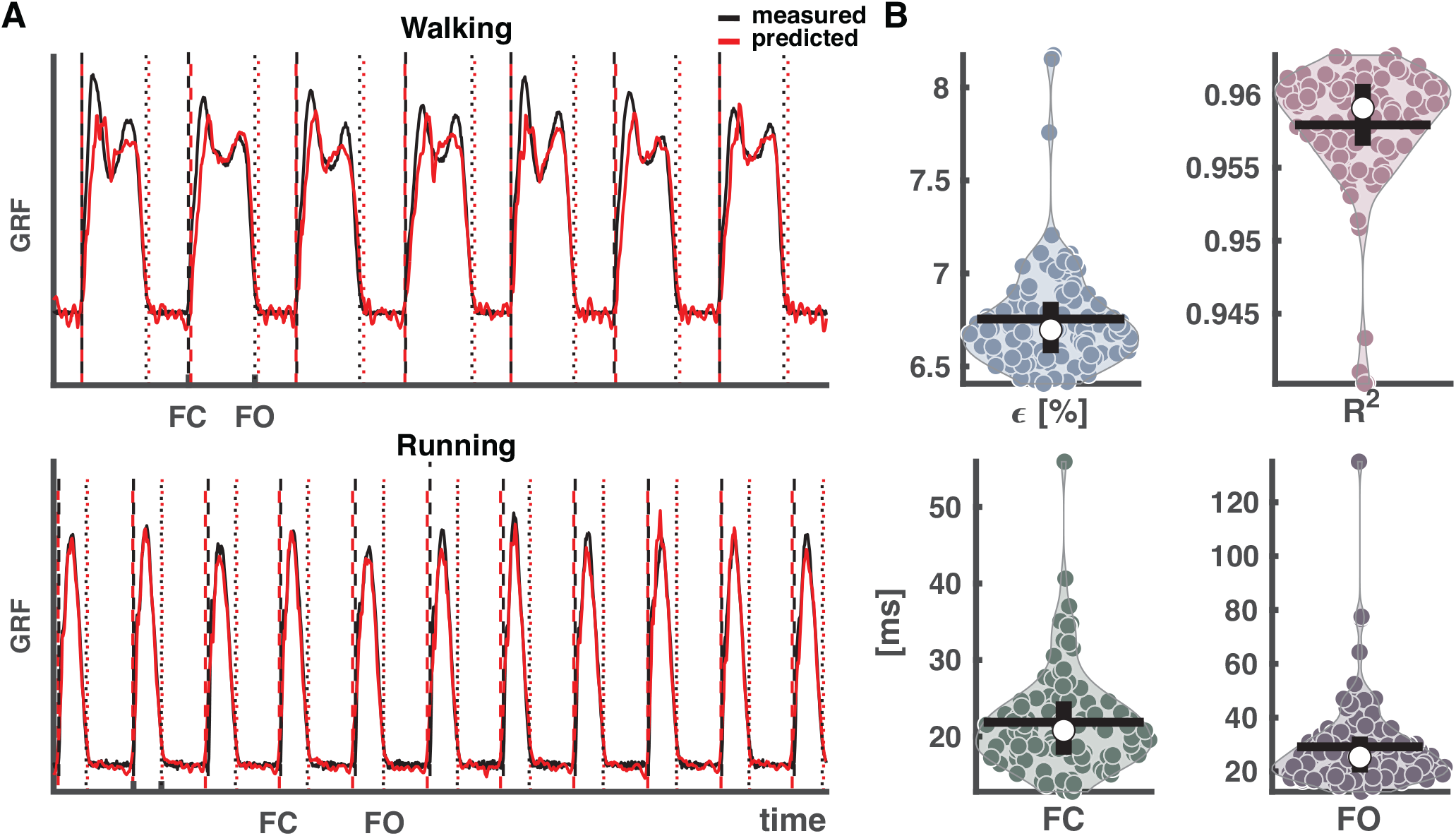
Output of reservoir computer trained on segmented data and validated and tested on continuous data pooled over conditions. (A) Vertical ground reaction force (GRF) waveforms of four randomly selected strides from each condition from a random trial and a random selected training run out of the 100, with the measured vertical GRF waveforms in black and predicted in red. The vertical dashed lines represent the foot contact events (black: measured, red: predicted), the vertical dotted lines the foot off events (black: measured, red: predicted). (B) Normalized root-mean squared error (***ϵ***), coefficient of determination (***R***^**2**^), mean absolute error of foot contact and foot off. The white dots in the violin-plots illustrate the medians. Black horizontal lines represent the mean and vertical black lines the 1st and 3rd quartiles. Every dot represents one of the 100 training runs, and the width of the violins is determined by the frequency. Abbreviations: GRF: ground reaction forces, FC: foot contact, FO: foot off.

To estimate the smallest number of strides needed for a mean reconstruction accuracy above 95%, we changed the size of the training set from 4 to 50% for the total dataset size (again with maximum 25 strides per trial for the training); cf. Figure 5. The size of the validation and test sets were kept fixed at 25% each to guarantee identical accuracy demands. An average of approximately 222 strides, ∼17% of the total dataset sufficed to reach *R*^2^ = 0.95 ± 0.01 with *ϵ* = 7.2 ± 0.3% and a mean absolute error of the foot contact (foot off) of 26.4 ± 9.3 ms (35.8 ± 15.8 ms).

**Figure 5:**
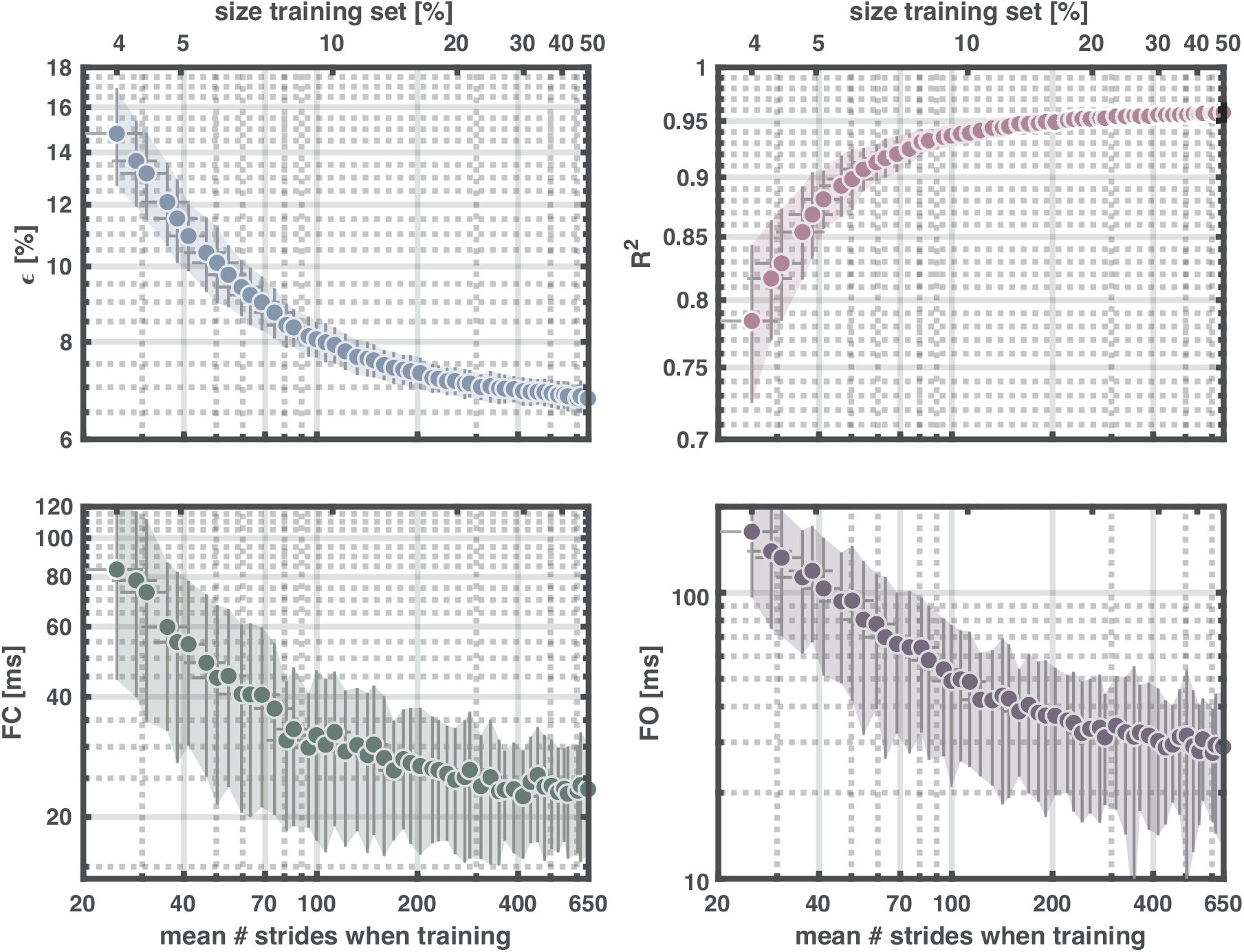
Output of reservoir computer with training data ranging from 4-50% of the total dataset and validating and testing with 25%, respectively. Upper left panel: normalized root-mean squared error (***ϵ***), upper right panel: coefficient of determination (***R***^**2**^), lower left panel: mean absolute error of foot contact, and lower right panel: mean absolute error of foot off. Upper x-axes show the percentage of the total dataset used for training and the lower x-axes the corresponding mean number of strides. Each dot represents the mean value for 100 repetitions. The vertical error-bars and colored areas represent the standard deviation of the corresponding measure. The horizontal error-bars represent the standard deviation in the number of strides across the 100 repetitions. Abbreviations: FC: foot contact, FO: foot off.

Finally, to test whether our approach allows for predicting vertical GRFs in trials that in their entirety were not part of the learning set, we performed a leave-M-out cross-validation. Training was realized in the held in trials using a 75%-25% split of learning and validation (in the training set we used a maximum number of 25 strides per trial). For *M* = 1, the mean *R*^2^ reached 0.91 ± 0.12, with an error *ϵ* of 9.1 ± 3.6%. One trial was a clear outlier in the leave-one-out analysis, and when re-calculating the means without this trial, the mean *R*^2^ and the error *ϵ* were 0.93 ± 0.04 and 8.6 ± 2.3%, respectively. The corresponding mean absolute errors for foot contact and foot off were 63.7 ± 167.1 ms and 140.9 ± 224.3 ms for all included trials and 49.0±138.9 ms and 129.8 ± 215.1 ms when not taking the outlier trial into account. For an overview of the results of *M* = 1 to *M* = 6, we refer to Supplemental Fig. S2.

## 4 Discussion

Reservoir computers are a promising tool to predict vertical GRF waveforms based on accelerometer data measured at the shank during walking and running. Accurately predicted GRF waveforms facilitate the detection of gait phases and events. We showed that this ‘simple’ machine learning approach has excellent prediction accuracy of continuous vertical GRF waveforms independent of the type of locomotion. Put differently, reservoir computers can be used for predicting vertical GRF waveforms for gait of unknown type with excellent performance. This has great potential for uses outside the lab and for collecting large amounts of data. Without a doubt the growing amount of data available for biomechanical analysis in running will greatly drive the field forwards (76). Machine learning combined with wearable sensors may be the solution to increase the amount of data recorded.

Using machine learning for activity recognition and gait phase recognition based on gait features extracted from biomechanical data, which may be measured with wearable sensors (77, 78) has become increasingly popular. The most common techniques to classify gait events or predict GRF waveforms using machine learning are hidden Markov models (26, 30, 33, 34, 37, 43, 79-82), neural networks such as deep neural networks (more than 1 hidden layer) (25, 35, 36, 44, 47, 52); feed-forward neural networks (48, 50, 53, 56-58, 83, 84); long short-term models (24, 28, 38, 54, 83, 84); convolutional neural networks (29, 55); support vector machines (42, 44, 85); (multilayer) perceptron models (28, 42, 49, 51, 86), as well as random forest classifiers (36, 42, 44), K-nearest neighbors (42, 54, 87), and other types of machine learning using, e.g., Bayesian models (31, 32, 82, 85), Gaussian mixture model (41), and principal component analysis (39, 40, 51, 82). Reservoir computers have the great advantage of low computational costs while still showing excellent performance. They merely require a handful of time series for training and avoid any a priori feature extraction. Reservoir computers even seem to be promising as a tool to successfully reproduce locomotor patterns observed during walking and running (88).

There have been several studies conducted on predicting GRFs based on wearables, the vast majority for the stance phase only. Utilizing these machine learning techniques properly requires foot contact and foot off to be known. This, however, can only be accomplished by either co-registering footswitches or ground reaction forces, or by implementing a rule-based detection of the gait events. This is exactly what we sought to circumvent. Rule-based detections based on accelerometers during running, searching for a specific peak or valley in some area of interest, are not as precise as an event detection based on vertical ground reaction forces.

We did not restrict the prediction to only the stance phase – we included the entire gait cycle and showed that this worked well on continuous data. We also did not time-normalize the gait cycles and did not impose any other constraints into the timing of the signals. As such, our predictions are robust against variations in speed and stride durations/lengths as well as the type of locomotion. For training the reservoir computer, data were segmented based on the ground truth events, though when these are unknown, the data could be segmented in any way, or they may not be segmented at all; cf. Figure 4. We are convinced that our approach is suitable for lab as well as outdoor use. One very recent study (89) predicted continuous vertical GRFs from trunk accelerations using a long short-term model network with good accuracy during sloped running. The pre-processing involved several filtering and feature extraction steps. By contrast, here we succeeded to reduce the number of pre-processing steps and applied only very weak filtering, i.e., the inputs are by and large the time series (derived from) vertical acceleration.

We validated the use of accelerometer data to estimate the vertical GRF waveform. We used accelerometer data collected by a sensor that could, in fact, also be used for electromyography data collection. The sensors were not aligned in the same way for all participants, which Tan, Chiasson (90) found can negatively affect the precision of the detection of GRFs using machine learning. However, variability in orientation and position is likely to occur if participants mount their own devices or in large-scale studies. By correcting the orientation of the accelerometers using principal component analysis we circumvent these potential problems and underline the robustness of our method and its applicability in many settings.

Even a small number of strides sufficed to achieve a high reconstruction accuracy. While strides from all participants were pooled for this analysis, it is unlikely that the strides randomly drawn into the training sets were representative for all trials/conditions and that this was also the case for the test sets. We evaluate this via leave-M-out cross-validation, as the test-set should be unknown to the machine (70). Admittedly, the reconstruction accuracy was not as high, but we consider it still satisfactory.

We would like to note that we did not optimize the machine learner to perform the absolute best it can do. Our primary aim was to show that even in its ‘simplest’ form, a reservoir computer with ridge-regression-based output weights can perform well. Apparently, this (off-line) approach has its limits as, during learning, one must store all network states which can put pressure on computer memory. The alternative online learning may be realized via recursive least squares regression (91), that has recently be adopted by Sussillo and Abbott (92). Along these lines one may add online feedback and change the reservoir’s connectivity for the network’s dynamics to reach the chaotic regime (currently we used a spectral radius of 0.5 but values larger than 1 may accelerate online learning; 92, 93). For our proof-of-concept, however, fine-tuning the reservoir might be considered overfitting, which let us decide not to progress along this direction. The most optimal settings will probably depend on the data set under study.

Our accelerometers had a relatively low sampling rate (≈ 143 Hz), which prevents better estimation than 7 ms (i.e., one frame equals 7 ms). An accelerometer with higher sampling frequency will arguably lead to higher accuracy of the predicted events compared to the ground truth. A sampling frequency of 60-200 Hz is not uncommon when recording kinematics (94-97) and the accuracy is not worse than the accuracy one could obtain using kinematic data. A frequently employed detection algorithm for kinematic gait event detection is the coordinate-based detection algorithm where the distance between the sacrum and foot is used to predict foot contact and foot off events (96, currently cited >800 times). A review on this and other detection algorithms (97) during running revealed that the coordinate-based detection algorithm has an absolute error of 29 ms for foot contact and 98 ms for foot off (sampling frequency: 200 Hz) whereas the best performing algorithms has an accuracy of 24 ms for foot contact (the foot vertical position (98, >600 citations)) and 6 ms for foot off (the peak knee extension algorithm (99, >600 citations)). For comparison, the best estimation possible with a sampling frequency of 200 Hz is 5 ms, i.e., current algorithms have an accuracy between one and ∼ 20 samples. Our approach is comparable to or exceeding this accuracy. Being cheap and easy to collect, being usable outside the lab and for long time-periods are, hence, not the only advantages of accelerometers – they also come with formidable accuracy in step detection when properly combined with reservoir computers.

Our machine learning approach performs well for a dataset comprised of both walking and running data despite a relatively small number of participants and a relatively small number of strides. To investigate its ability for each condition separately, we refer to Supplemental Fig. S1, where we show that training and testing on only one type of locomotion improves the already excellent reconstruction accuracy. The outputs of the reservoir computer can easily be modified to provide other outputs such as other components of the GRF or the center-of-pressure. By including all three components of the GRF, energetics of the center-of-mass can be estimated during overground/track running which in turn can provide even more information about the locomotion type. Of course, the energetics will be in arbitrary units given our GRF prediction rely on normalize (z-scored) values. All data for this study were recorded on the treadmill. The next step will be to apply reservoir-based prediction to accelerometer data (or gyroscope data) obtained at other parts of the body, e.g., hip mounted (e.g., activity trackers), arm mounted (e.g., sport watches, smartphones) or head mounted (e.g., augmented/virtual reality glasses) to broaden applicability in daily living contexts as well as in clinical populations as machine learning algorithms might perform worse on clinical gait (100). This certainly calls for expanding the current dataset with overground/outdoor locomotion. We expect our findings to be transferable to overground settings. A large meta-analysis suggests that neither vertical ground reaction forces, nor peak tibial accelerations are significantly different between treadmill and overground running (101). However, this might not be true when accelerations, decelerations, turns, etc. are considered.

As a final note we would like to recall that our reservoir computer did not include a feedback loop. Adding feedback may allow for not only predicting the GRF accompanying tibial accelerations but eventually also the GRF in forthcoming strides. We trust that future studies will pursue this generalization as it is beyond the scope of our proof-of-concept study. Given our formidable prediction results, however, we can stress that all the information needed for predicting (vertical) ground reaction forces seems to be present in the (principal component of) accelerometer signals. The use of the latter, hence, provides more opportunities than commonly thought.

Our data and code are made freely available and are ready to use on other datasets and can be extended for use in experiments and clinic.

## 5 Conclusion

Reservoir computers are an excellent candidate to correctly predict vertical ground reaction force waveforms from accelerometer signals for a small number of participants and strides. The predicted time series can serve to estimate stepping moments with particularly high accuracy. The ease in training procedure, which requires only a (very) limited number of steps and without prior knowledge about the type of locomotion lets us advocate this machine learning approach to be further expanded to be applied on future applications in both research and clinic.

## 7 Author Contributions

M.B. and N.D. set up the experiment, M.B. collected and pre-processed the data, M.B. and A.D. wrote the code, M.B. prepared all figures, M.B. and A.D. wrote the first draft of the manuscript. All authors commented and provided feedback on the final version of the manuscript.

## 9 List of abbreviations

GRF: Ground reaction forces
ESN: Echo state network
PCA: Principal component analysis

## 10 Acknowledgments

We would like to acknowledge the participants who participated in the study and Emiel den Haan and Rohan Zonneveld for the data collection. We would also like to acknowledge Sjoerd Bruijn and Jaap van Dieën for their valuable inputs to a previous version of the manuscript.

## 11 Data Availability Statement

The datasets analyzed and the code produced during the current study are available in the GitHub repository, https://github.com/marlow17/PredictingGroundReactionForces.

## References

1. van Oeveren BT, de Ruiter CJ, Beek PJ, Rispens SM, van Dieen JH. An Adaptive, Real-Time Cadence Algorithm for Unconstrained Sensor Placement. Med Eng Phys (2018) 52:49–58. Epub 2018/01/27. doi: 10.1016/j.medengphy.2017.12.007.

2. van Oeveren B. Running Deciphered: The Interpretation of Running Technique from Wearable Data. Amsterdam: Vrije Universiteit Amsterdam (2021).

3. de Ruiter CJ, van Oeveren B, Francke A, Zijlstra P, van Dieen JH. Running Speed Can Be Predicted from Foot Contact Time During Outdoor over Ground Running. PLoS One (2016) 11(9):e0163023. Epub 2016/09/21. doi: 10.1371/journal.pone.0163023.

4. Moe-Nilssen R, Helbostad JL. Estimation of Gait Cycle Characteristics by Trunk Accelerometry. J Biomech (2004) 37(1):121–6. Epub 2003/12/16. doi: 10.1016/s0021-9290(03)00233-1.

5. Norris M, Kenny IC, Anderson R. Comparison of Accelerometry Stride Time Calculation Methods. J Biomech (2016) 49(13):3031–4. Epub 2016/06/13. doi: 10.1016/j.jbiomech.2016.05.029.

6. Pérez-Ibarra JC, Williams H, Siqueira AA, Krebs HI, editors. Real-Time Identification of Impaired Gait Phases Using a Single Foot-Mounted Inertial Sensor: Review and Feasibility Study. 2018 7th IEEE International Conference on Biomedical Robotics and Biomechatronics (Biorob); 2018: IEEE.

7. Prasanth H, Caban M, Keller U, Courtine G, Ijspeert A, Vallery H, et al. Wearable Sensor-Based Real-Time Gait Detection: A Systematic Review. Sensors (Basel) (2021) 21(8). Epub 2021/05/01. doi: 10.3390/s21082727.

8. Trojaniello D, Cereatti A, Pelosin E, Avanzino L, Mirelman A, Hausdorff JM, et al. Estimation of Step-by-Step Spatio-Temporal Parameters of Normal and Impaired Gait Using Shank-Mounted Magneto-Inertial Sensors: Application to Elderly, Hemiparetic, Parkinsonian and Choreic Gait. J Neuroeng Rehabil (2014) 11:152. Epub 2014/11/13. doi: 10.1186/1743-0003-11-152.

9. Rueterbories J, Spaich EG, Andersen OK. Gait Event Detection for Use in Fes Rehabilitation by Radial and Tangential Foot Accelerations. Med Eng Phys (2014) 36(4):502–8. Epub 2013/11/05. doi: 10.1016/j.medengphy.2013.10.004.

10. Ben Mansour K, Rezzoug N, Gorce P. Analysis of Several Methods and Inertial Sensors Locations to Assess Gait Parameters in Able-Bodied Subjects. Gait Posture (2015) 42(4):409–14. Epub 2015/09/06. doi: 10.1016/j.gaitpost.2015.05.020.

11. Greene BR, McGrath D, O’Neill R, O’Donovan KJ, Burns A, Caulfield B. An Adaptive Gyroscope-Based Algorithm for Temporal Gait Analysis. Med Biol Eng Comput (2010) 48(12):1251–60. Epub 2010/11/03. doi: 10.1007/s11517-010-0692-0.

12. Pacini Panebianco G, Bisi MC, Stagni R, Fantozzi S. Analysis of the Performance of 17 Algorithms from a Systematic Review: Influence of Sensor Position, Analysed Variable and Computational Approach in Gait Timing Estimation from Imu Measurements. Gait Posture (2018) 66:76–82. Epub 2018/09/01. doi: 10.1016/j.gaitpost.2018.08.025.

13. Selles RW, Formanoy MA, Bussmann JB, Janssens PJ, Stam HJ. Automated Estimation of Initial and Terminal Contact Timing Using Accelerometers; Development and Validation in Transtibial Amputees and Controls. IEEE Trans Neural Syst Rehabil Eng (2005) 13(1):81–8. Epub 2005/04/09. doi: 10.1109/TNSRE.2004.843176.

14. Mico-Amigo ME, Kingma I, Ainsworth E, Walgaard S, Niessen M, van Lummel RC, et al. A Novel Accelerometry-Based Algorithm for the Detection of Step Durations over Short Episodes of Gait in Healthy Elderly. J Neuroeng Rehabil (2016) 13:38. Epub 2016/04/21. doi: 10.1186/s12984-016-0145-6.

15. Gurchiek RD, Garabed CP, McGinnis RS. Gait Event Detection Using a Thigh-Worn Accelerometer. Gait Posture (2020) 80:214–6. Epub 2020/06/15. doi: 10.1016/j.gaitpost.2020.06.004.

16. Mo S, Chow DHK. Accuracy of Three Methods in Gait Event Detection During Overground Running. Gait Posture (2018) 59:93–8. Epub 2017/10/14. doi: 10.1016/j.gaitpost.2017.10.009.

17. Khandelwal S, Wickstrom N. Evaluation of the Performance of Accelerometer-Based Gait Event Detection Algorithms in Different Real-World Scenarios Using the Marea Gait Database. Gait Posture (2017) 51:84–90. Epub 2016/10/14. doi: 10.1016/j.gaitpost.2016.09.023.

18. Mitschke C, Heß T, Milani T. Which Method Detects Foot Strike in Rearfoot and Forefoot Runners Accurately When Using an Inertial Measurement Unit? Applied Sciences (2017) 7(9):959.

19. Sinclair J, Hobbs SJ, Protheroe L, Edmundson CJ, Greenhalgh A. Determination of Gait Events Using an Externally Mounted Shank Accelerometer. J Appl Biomech (2013) 29(1):118–22. Epub 2013/03/07. doi: 10.1123/jab.29.1.118.

20. Purcell B, Channells J, James D, Barrett R, editors. Use of Accelerometers for Detecting Foot-Ground Contact Time During Running. BioMEMS and Nanotechnology II; 2006: International Society for Optics and Photonics.

21. Lee JB, Mellifont RB, Burkett BJ. The Use of a Single Inertial Sensor to Identify Stride, Step, and Stance Durations of Running Gait. Journal of Science and Medicine in Sport (2010) 13(2):270–3.

22. McGrath D, Greene BR, O’Donovan KJ, Caulfield B. Gyroscope-Based Assessment of Temporal Gait Parameters During Treadmill Walking and Running. Sports Engineering (2012) 15(4):207–13. doi: 10.1007/s12283-012-0093-8.

23. Bergamini E, Picerno P, Pillet H, Natta F, Thoreux P, Camomilla V. Estimation of Temporal Parameters During Sprint Running Using a Trunk-Mounted Inertial Measurement Unit. J Biomech (2012) 45(6):1123–6. Epub 2012/02/14. doi: 10.1016/j.jbiomech.2011.12.020.

24. Tan HX, Aung NN, Tian J, Chua MCH, Yang YO. Time Series Classification Using a Modified Lstm Approach from Accelerometer-Based Data: A Comparative Study for Gait Cycle Detection. Gait Posture (2019) 74:128–34. Epub 2019/09/14. doi: 10.1016/j.gaitpost.2019.09.007.

25. Prado A, Cao X, Robert MT, Gordon AM, Agrawal SK. Gait Segmentation of Data Collected by Instrumented Shoes Using a Recurrent Neural Network Classifier. Phys Med Rehabil Clin N Am (2019) 30(2):355–66. Epub 2019/04/08. doi: 10.1016/j.pmr.2018.12.007.

26. Mannini A, Genovese V, Maria Sabatini A. Online Decoding of Hidden Markov Models for Gait Event Detection Using Foot-Mounted Gyroscopes. IEEE J Biomed Health Inform (2014) 18(4):1122–30. Epub 2014/07/12. doi: 10.1109/JBHI.2013.2293887.

27. Mannini A, Sabatini AM, editors. A Hidden Markov Model-Based Technique for Gait Segmentation Using a Foot-Mounted Gyroscope. 2011 Annual International Conference of the IEEE Engineering in Medicine and Biology Society; 2011: IEEE.

28. Robberechts P, Derie R, Van den Berghe P, Gerlo J, De Clercq D, Segers V, et al. Predicting Gait Events from Tibial Acceleration in Rearfoot Running: A Structured Machine Learning Approach. Gait Posture (2021) 84:87–92. Epub 2020/12/08. doi: 10.1016/j.gaitpost.2020.10.035.

29. Su B, Smith C, Gutierrez Farewik E. Gait Phase Recognition Using Deep Convolutional Neural Network with Inertial Measurement Units. Biosensors (Basel) (2020) 10(9). Epub 2020/09/02. doi: 10.3390/bios10090109.

30. Taborri J, Rossi S, Palermo E, Patane F, Cappa P. A Novel Hmm Distributed Classifier for the Detection of Gait Phases by Means of a Wearable Inertial Sensor Network. Sensors (Basel) (2014) 14(9):16212–34. Epub 2014/09/04. doi: 10.3390/s140916212.

31. Martinez-Hernandez U, Dehghani-Sanij AA. Adaptive Bayesian Inference System for Recognition of Walking Activities and Prediction of Gait Events Using Wearable Sensors. Neural Netw (2018) 102:107–19. Epub 2018/03/24. doi: 10.1016/j.neunet.2018.02.017.

32. Meng L, Martinez-Hernandez U, Childs C, Dehghani-Sanij AA, Buis A. A Practical Gait Feedback Method Based on Wearable Inertial Sensors for a Drop Foot Assistance Device. IEEE Sensors Journal (2019) 19(24):12235–43. doi: 10.1109/jsen.2019.2938764.

33. Taborri J, Scalona E, Palermo E, Rossi S, Cappa P. Validation of Inter-Subject Training for Hidden Markov Models Applied to Gait Phase Detection in Children with Cerebral Palsy. Sensors (Basel) (2015) 15(9):24514–29. Epub 2015/09/26. doi: 10.3390/s150924514.

34. Abaid N, Cappa P, Palermo E, Petrarca M, Porfiri M. Gait Detection in Children with and without Hemiplegia Using Single-Axis Wearable Gyroscopes. PLoS One (2013) 8(9):e73152. Epub 2013/09/12. doi: 10.1371/journal.pone.0073152.

35. Vu HTT, Gomez F, Cherelle P, Lefeber D, Nowe A, Vanderborght B. Ed-Fnn: A New Deep Learning Algorithm to Detect Percentage of the Gait Cycle for Powered Prostheses. Sensors (Basel) (2018) 18(7). Epub 2018/07/26. doi: 10.3390/s18072389.

36. Yang J, Huang T-H, Yu S, Yang X, Su H, Spungen AM, et al., editors. Machine Learning Based Adaptive Gait Phase Estimation Using Inertial Measurement Sensors. Frontiers in Biomedical Devices; 2019: American Society of Mechanical Engineers.

37. Chen G, Qi P, Guo Z, Yu H. Gait-Event-Based Synchronization Method for Gait Rehabilitation Robots Via a Bioinspired Adaptive Oscillator. IEEE Trans Biomed Eng (2017) 64(6):1345–56. Epub 2017/01/24. doi: 10.1109/TBME.2016.2604340.

38. Kidzinski L, Delp S, Schwartz M. Automatic Real-Time Gait Event Detection in Children Using Deep Neural Networks. PLoS One (2019) 14(1):e0211466. Epub 2019/02/01. doi: 10.1371/journal.pone.0211466.

39. Osis ST, Hettinga BA, Ferber R. Predicting Ground Contact Events for a Continuum of Gait Types: An Application of Targeted Machine Learning Using Principal Component Analysis. Gait Posture (2016) 46:86–90. Epub 2016/05/01. doi: 10.1016/j.gaitpost.2016.02.021.

40. Osis ST, Hettinga BA, Leitch J, Ferber R. Predicting Timing of Foot Strike During Running, Independent of Striking Technique, Using Principal Component Analysis of Joint Angles. J Biomech (2014) 47(11):2786–9. Epub 2014/07/12. doi: 10.1016/j.jbiomech.2014.06.009.

41. Aung MS, Thies SB, Kenney LP, Howard D, Selles RW, Findlow AH, et al. Automated Detection of Instantaneous Gait Events Using Time Frequency Analysis and Manifold Embedding. IEEE Trans Neural Syst Rehabil Eng (2013) 21(6):908–16. Epub 2013/01/17. doi: 10.1109/TNSRE.2013.2239313.

42. Morbidoni C, Cucchiarelli A, Agostini V, Knaflitz M, Fioretti S, Di Nardo F. Machine-Learning-Based Prediction of Gait Events from Emg in Cerebral Palsy Children. IEEE Trans Neural Syst Rehabil Eng (2021) 29:819–30. Epub 2021/04/29. doi: 10.1109/TNSRE.2021.3076366.

43. Crea S, De Rossi SM, Donati M, Rebersek P, Novak D, Vitiello N, et al. Development of Gait Segmentation Methods for Wearable Foot Pressure Sensors. Annu Int Conf IEEE Eng Med Biol Soc (2012) 2012:5018–21. Epub 2013/02/01. doi: 10.1109/EMBC.2012.6347120.

44. Rezaei A, Ejupi A, Gholami M, Ferrone A, Menon C, editors. Preliminary Investigation of Textile-Based Strain Sensors for the Detection of Human Gait Phases Using Machine Learning. 2018 7th IEEE International Conference on Biomedical Robotics and Biomechatronics (Biorob); 2018: IEEE.

45. Roerdink M, Coolen BH, Clairbois BH, Lamoth CJ, Beek PJ. Online Gait Event Detection Using a Large Force Platform Embedded in a Treadmill. J Biomech (2008) 41(12):2628–32. Epub 2008/07/29. doi: 10.1016/j.jbiomech.2008.06.023.

46. Borghese NA, Bianchi L, Lacquaniti F. Kinematic Determinants of Human Locomotion. J Physiol (1996) 494 (Pt 3):863–79. Epub 1996/08/01. doi: 10.1113/jphysiol.1996.sp021539.

47. Davidson P, Virekunnas H, Sharma D, Piche R, Cronin N. Continuous Analysis of Running Mechanics by Means of an Integrated Ins/Gps Device. Sensors (Basel) (2019) 19(6). Epub 2019/03/29. doi: 10.3390/s19061480.

48. Ngoh KJ, Gouwanda D, Gopalai AA, Chong YZ. Estimation of Vertical Ground Reaction Force During Running Using Neural Network Model and Uniaxial Accelerometer. J Biomech (2018) 76:269–73. Epub 2018/06/28. doi: 10.1016/j.jbiomech.2018.06.006.

49. Leporace G, Batista LA, Nadal J. Prediction of 3d Ground Reaction Forces During Gait Based on Accelerometer Data. Research on Biomedical Engineering (2018) 34(3):211–6. doi: 10.1590/2446-4740.06817.

50. Lim H, Kim B, Park S. Prediction of Lower Limb Kinetics and Kinematics During Walking by a Single Imu on the Lower Back Using Machine Learning. Sensors (Basel) (2019) 20(1). Epub 2019/12/28. doi: 10.3390/s20010130.

51. Pogson M, Verheul J, Robinson MA, Vanrenterghem J, Lisboa P. A Neural Network Method to Predict Task-and Step-Specific Ground Reaction Force Magnitudes from Trunk Accelerations During Running Activities. Med Eng Phys (2020) 78:82–9. Epub 2020/03/03. doi: 10.1016/j.medengphy.2020.02.002.

52. Wouda FJ, Giuberti M, Bellusci G, Maartens E, Reenalda J, van Beijnum BF, et al. Estimation of Vertical Ground Reaction Forces and Sagittal Knee Kinematics During Running Using Three Inertial Sensors. Front Physiol (2018) 9:218. Epub 2018/04/07. doi: 10.3389/fphys.2018.00218.

53. Lee M, Park S. Estimation of Three-Dimensional Lower Limb Kinetics Data During Walking Using Machine Learning from a Single Imu Attached to the Sacrum. Sensors (Basel) (2020) 20(21). Epub 2020/11/08. doi: 10.3390/s20216277.

54. Sharma D, Davidson P, Muller P, Piche R. Indirect Estimation of Vertical Ground Reaction Force from a Body-Mounted Ins/Gps Using Machine Learning. Sensors (Basel) (2021) 21(4). Epub 2021/03/07. doi: 10.3390/s21041553.

55. Johnson WR, Mian A, Robinson MA, Verheul J, Lloyd DG, Alderson JA. Multidimensional Ground Reaction Forces and Moments from Wearable Sensor Accelerations Via Deep Learning. IEEE Trans Biomed Eng (2021) 68(1):289–97. Epub 2020/08/04. doi: 10.1109/TBME.2020.3006158.

56. Komaris D-S, Perez-Valero E, Jordan L, Barton J, Hennessy L, O’Flynn B, et al. Predicting Three-Dimensional Ground Reaction Forces in Running by Using Artificial Neural Networks and Lower Body Kinematics. IEEE Access (2019) 7:156779–86. doi: 10.1109/access.2019.2949699.

57. Choi A, Lee J-M, Mun JH. Ground Reaction Forces Predicted by Using Artificial Neural Network During Asymmetric Movements. International Journal of Precision Engineering and Manufacturing (2013) 14(3):475–83. doi: 10.1007/s12541-013-0064-4.

58. Oh SE, Choi A, Mun JH. Prediction of Ground Reaction Forces During Gait Based on Kinematics and a Neural Network Model. J Biomech (2013) 46(14):2372–80. Epub 2013/08/22. doi: 10.1016/j.jbiomech.2013.07.036.

59. Guo Y, Storm F, Zhao Y, Billings SA, Pavic A, Mazza C, et al. A New Proxy Measurement Algorithm with Application to the Estimation of Vertical Ground Reaction Forces Using Wearable Sensors. Sensors (Basel) (2017) 17(10). Epub 2017/09/25. doi: 10.3390/s17102181.

60. Horsley BJ, Tofari PJ, Halson SL, Kemp JG, Dickson J, Maniar N, et al. Does Site Matter? Impact of Inertial Measurement Unit Placement on the Validity and Reliability of Stride Variables During Running: A Systematic Review and Meta-Analysis. Sports Med (2021) 51(7):1449–89. Epub 2021/03/25. doi: 10.1007/s40279-021-01443-8.

61. Ancillao A, Tedesco S, Barton J, O’Flynn B. Indirect Measurement of Ground Reaction Forces and Moments by Means of Wearable Inertial Sensors: A Systematic Review. Sensors (Basel) (2018) 18(8). Epub 2018/08/08. doi: 10.3390/s18082564.

62. Jaeger H, Haas H. Harnessing Nonlinearity: Predicting Chaotic Systems and Saving Energy in Wireless Communication. Science (2004) 304(5667):78–80. Epub 2004/04/06. doi: 10.1126/science.1091277.

63. Goodfellow I, Bengio Y, Courville A. Deep Learning: MIT press (2016).

64. Maass W, Natschläger T, Markram H. Real-Time Computing without Stable States: A New Framework for Neural Computation Based on Perturbations. Neural computation (2002) 14(11):2531–60.

65. Pathak J, Hunt B, Girvan M, Lu Z, Ott E. Model-Free Prediction of Large Spatiotemporally Chaotic Systems from Data: A Reservoir Computing Approach. Phys Rev Lett (2018) 120(2):024102. Epub 2018/01/30. doi: 10.1103/PhysRevLett.120.024102.

66. Lukoševičius M, Jaeger H. Reservoir Computing Approaches to Recurrent Neural Network Training. Computer Science Review (2009) 3(3):127–49. doi: 10.1016/j.cosrev.2009.03.005.

67. Jordan K, Challis JH, Newell KM. Walking Speed Influences on Gait Cycle Variability. Gait Posture (2007) 26(1):128–34. Epub 2006/09/20. doi: 10.1016/j.gaitpost.2006.08.010.

68. Moe-Nilssen R. A New Method for Evaluating Motor Control in Gait under Real-Life Environmental Conditions. Part 1: The Instrument. Clin Biomech (Bristol, Avon) (1998) 13(4-5):320–7. Epub 2001/06/21. doi: 10.1016/s0268-0033(98)00089-8.

69. Rispens SM, Pijnappels M, van Schooten KS, Beek PJ, Daffertshofer A, van Dieen JH. Consistency of Gait Characteristics as Determined from Acceleration Data Collected at Different Trunk Locations. Gait Posture (2014) 40(1):187–92. Epub 2014/05/02. doi: 10.1016/j.gaitpost.2014.03.182.

70. Halilaj E, Rajagopal A, Fiterau M, Hicks JL, Hastie TJ, Delp SL. Machine Learning in Human Movement Biomechanics: Best Practices, Common Pitfalls, and New Opportunities. J Biomech (2018) 81:1–11. Epub 2018/10/04. doi: 10.1016/j.jbiomech.2018.09.009.

71. Savitzky A, Golay MJ. Smoothing and Differentiation of Data by Simplified Least Squares Procedures. Analytical chemistry (1964) 36(8):1627–39.

72. Ghoussayni S, Stevens C, Durham S, Ewins D. Assessment and Validation of a Simple Automated Method for the Detection of Gait Events and Intervals. Gait Posture (2004) 20(3):266–72. Epub 2004/11/09. doi: 10.1016/j.gaitpost.2003.10.001.

73. Jaeger H. The “Echo State” Approach to Analysing and Training Recurrent Neural Networks-with an Erratum Note. Bonn, Germany: German National Research Center for Information Technology GMD Technical Report (2001) 148(34):13.

74. Jaeger H. Tutorial on Training Recurrent Neural Networks, Covering Bppt, Rtrl, Ekf and the “Echo State Network” Approach: GMD-Forschungszentrum Informationstechnik Bonn (2002).

75. Lukoševičius M. A Practical Guide to Applying Echo State Networks. In: Montavon G, Orr GB, Müller K-R, editors. Neural Networks: Tricks of the Trade: Second Edition. Berlin, Heidelberg: Springer Berlin Heidelberg (2012). p. 659–86.

76. Novacheck TF. The Biomechanics of Running. Gait Posture (1998) 7(1):77–95. Epub 1999/04/14. doi: 10.1016/s0966-6362(97)00038-6.

77. Farrahi V, Niemela M, Kangas M, Korpelainen R, Jamsa T. Calibration and Validation of Accelerometer-Based Activity Monitors: A Systematic Review of Machine-Learning Approaches. Gait Posture (2019) 68:285–99. Epub 2018/12/24. doi: 10.1016/j.gaitpost.2018.12.003.

78. Figueiredo J, Santos CP, Moreno JC. Automatic Recognition of Gait Patterns in Human Motor Disorders Using Machine Learning: A Review. Med Eng Phys (2018) 53:1–12. Epub 2018/01/27. doi: 10.1016/j.medengphy.2017.12.006.

79. Mannini A, Sabatini AM. Gait Phase Detection and Discrimination between Walking-Jogging Activities Using Hidden Markov Models Applied to Foot Motion Data from a Gyroscope. Gait Posture (2012) 36(4):657–61. Epub 2012/07/17. doi: 10.1016/j.gaitpost.2012.06.017.

80. Chen G, Salim V, Yu H, editors. A Novel Gait Phase-Based Control Strategy for a Portable Knee-Ankle-Foot Robot. 2015 IEEE International Conference on Rehabilitation Robotics (ICORR); 2015: IEEE.

81. Guenterberg E, Yang AY, Ghasemzadeh H, Jafari R, Bajcsy R, Sastry SS. A Method for Extracting Temporal Parameters Based on Hidden Markov Models in Body Sensor Networks with Inertial Sensors. IEEE Trans Inf Technol Biomed (2009) 13(6):1019–30. Epub 2009/09/04. doi: 10.1109/TITB.2009.2028421.

82. Yuwono M, Su SW, Guo Y, Moulton BD, Nguyen HT. Unsupervised Nonparametric Method for Gait Analysis Using a Waist-Worn Inertial Sensor. Applied Soft Computing (2014) 14:72–80. doi: 10.1016/j.asoc.2013.07.027.

83. Choi A, Jung H, Lee KY, Lee S, Mun JH. Machine Learning Approach to Predict Center of Pressure Trajectories in a Complete Gait Cycle: A Feedforward Neural Network Vs. Lstm Network. Med Biol Eng Comput (2019) 57(12):2693–703. Epub 2019/10/28. doi: 10.1007/s11517-019-02056-0.

84. Choi A, Jung H, Mun JH. Single Inertial Sensor-Based Neural Networks to Estimate Com-Cop Inclination Angle During Walking. Sensors (Basel) (2019) 19(13). Epub 2019/07/10. doi: 10.3390/s19132974.

85. Nutakki C, Mathew RJ, Suresh A, Vijay AR, Krishna S, Babu AS, et al. Classification and Kinetic Analysis of Healthy Gait Using Multiple Accelerometer Sensors. Procedia Computer Science (2020) 171:395–402.

86. Mijailovic N, Gavrilovic M, Rafajlovic S, Ðuric-Jovicic M, Popovic D. Gait Phases Recognition from Accelerations and Ground Reaction Forces: Application of Neural Networks. Telfor Journal (2009) 1(1):34–6.

87. Chu KH, Jiang X, Menon C. Wearable Step Counting Using a Force Myography-Based Ankle Strap. J Rehabil Assist Technol Eng (2017) 4:2055668317746307. Epub 2017/12/06. doi: 10.1177/2055668317746307.

88. de Graaf ML, Mochizuki L, Thies F, Wagner H, Le Mouel C. Motor Pattern Generation Is Robust to Neural Network Anatomical Imbalance Favoring Inhibition but Not Excitation. bioRxiv (2022):2022.04.21.489087. doi: 10.1101/2022.04.21.489087.

89. Alcantara RS, Edwards WB, Millet GY, Grabowski AM. Predicting Continuous Ground Reaction Forces from Accelerometers During Uphill and Downhill Running: A Recurrent Neural Network Solution. PeerJ (2022) 10:e12752. Epub 2022/01/18. doi: 10.7717/peerj.12752.

90. Tan T, Chiasson DP, Hu H, Shull PB. Influence of Imu Position and Orientation Placement Errors on Ground Reaction Force Estimation. J Biomech (2019) 97:109416. Epub 2019/10/22. doi: 10.1016/j.jbiomech.2019.109416.

91. Haykin S. Adaptive Filter Theory: International Edition: Pearson Education (2014).

92. Sussillo D, Abbott LF. Generating Coherent Patterns of Activity from Chaotic Neural Networks. Neuron (2009) 63(4):544–57. Epub 2009/08/28. doi: 10.1016/j.neuron.2009.07.018.

93. Yildiz IB, Jaeger H, Kiebel SJ. Re-Visiting the Echo State Property. Neural Netw (2012) 35:1–9. Epub 2012/08/14. doi: 10.1016/j.neunet.2012.07.005.

94. Hreljac A, Marshall RN. Algorithms to Determine Event Timing During Normal Walking Using Kinematic Data. J Biomech (2000) 33(6):783–6. Epub 2000/05/16. doi: 10.1016/s0021-9290(00)00014-2.

95. O’Connor CM, Thorpe SK, O’Malley MJ, Vaughan CL. Automatic Detection of Gait Events Using Kinematic Data. Gait Posture (2007) 25(3):469–74. Epub 2006/08/01. doi: 10.1016/j.gaitpost.2006.05.016.

96. Zeni JA, Jr., Richards JG, Higginson JS. Two Simple Methods for Determining Gait Events During Treadmill and Overground Walking Using Kinematic Data. Gait Posture (2008) 27(4):710–4. Epub 2007/08/29. doi: 10.1016/j.gaitpost.2007.07.007.

97. Fellin RE, Rose WC, Royer TD, Davis IS. Comparison of Methods for Kinematic Identification of Footstrike and Toe-Off During Overground and Treadmill Running. J Sci Med Sport (2010) 13(6):646–50. Epub 2010/05/19. doi: 10.1016/j.jsams.2010.03.006.

98. Alton F, Baldey L, Caplan S, Morrissey MC. A Kinematic Comparison of Overground and Treadmill Walking. Clin Biomech (Bristol, Avon) (1998) 13(6):434–40. Epub 2001/06/21. doi: 10.1016/s0268-0033(98)00012-6.

99. Dingwell JB, Cusumano JP, Cavanagh PR, Sternad D. Local Dynamic Stability Versus Kinematic Variability of Continuous Overground and Treadmill Walking. J Biomech Eng (2001) 123(1):27–32. Epub 2001/03/30. doi: 10.1115/1.1336798.

100. Bastien GJ, Gosseye TP, Penta M. A Robust Machine Learning Enabled Decomposition of Shear Ground Reaction Forces During the Double Contact Phase of Walking. Gait Posture (2019) 73:221–7. Epub 2019/08/03. doi: 10.1016/j.gaitpost.2019.07.190.

101. Van Hooren B, Fuller JT, Buckley JD, Miller JR, Sewell K, Rao G, et al. Is Motorized Treadmill Running Biomechanically Comparable to Overground Running? A Systematic Review and Meta-Analysis of Cross-over Studies. Sports Med (2020) 50(4):785–813. Epub 2019/12/06. doi: 10.1007/s40279-019-01237-z.

