## Supplemental Material for "Predicting vertical ground reaction forces from 3D accelerometry using reservoir computers leads to accurate gait event detection"

Predicting vertical ground reaction forces from 3D accelerometery using reservoir computers leads to accurate gait event detection, Margit M. Bach, Nadia Dominici*, and Andreas Daffertshofer*, Department of Human Movement Sciences, Faculty of Behavioural and Movement Sciences, Amsterdam Movement Sciences & Institute of Brain and Behavior Amsterdam, Vrije Universiteit Amsterdam, Amsterdam, The Netherlands

### *Supplementary Material*

To investigate if training the reservoir computer on each condition separately resulted in even better prediction values, we trained and validated on stride-segmented data and tested its prediction capacity for continuous gait data for each condition separately. For each trial, 25% of the continuous data was selected for testing, another 25% of the continuous data was selected for validation and the remaining 50% was segmented into strides (with a maximum of 25 strides per trial) and subsequently pooled across all trials and used for training totalling a 50/25/25% split for training, validation, and testing respectively. For both walking and running the quality of performance of predicting vertical GRF waveforms exceeded 97%: the coefficients of determination $R^2$ were $0.97 \pm 0.00$ and $0.97 \pm 0.02$, respectively, and the normalised root-mean squared errors $\epsilon$ were $5.9 \pm 0.4\%$ and $5.6 \pm 0.3\%$, respectively (mean ± SD); cf. Figure S1. On average, the subsequently extracted foot contact (foot off) events deviated from those based on the measured vertical GRF waveforms by 2.4 (2.5) samples for walking and 1.9 (2.3) samples for running. This corresponds to mean absolute errors of 17.3 ± 12.2 ms (17.5 ± 11.3 ms) for walking and 13.5 ± 3.9 ms (16.1 ± 8.5 ms) for running.

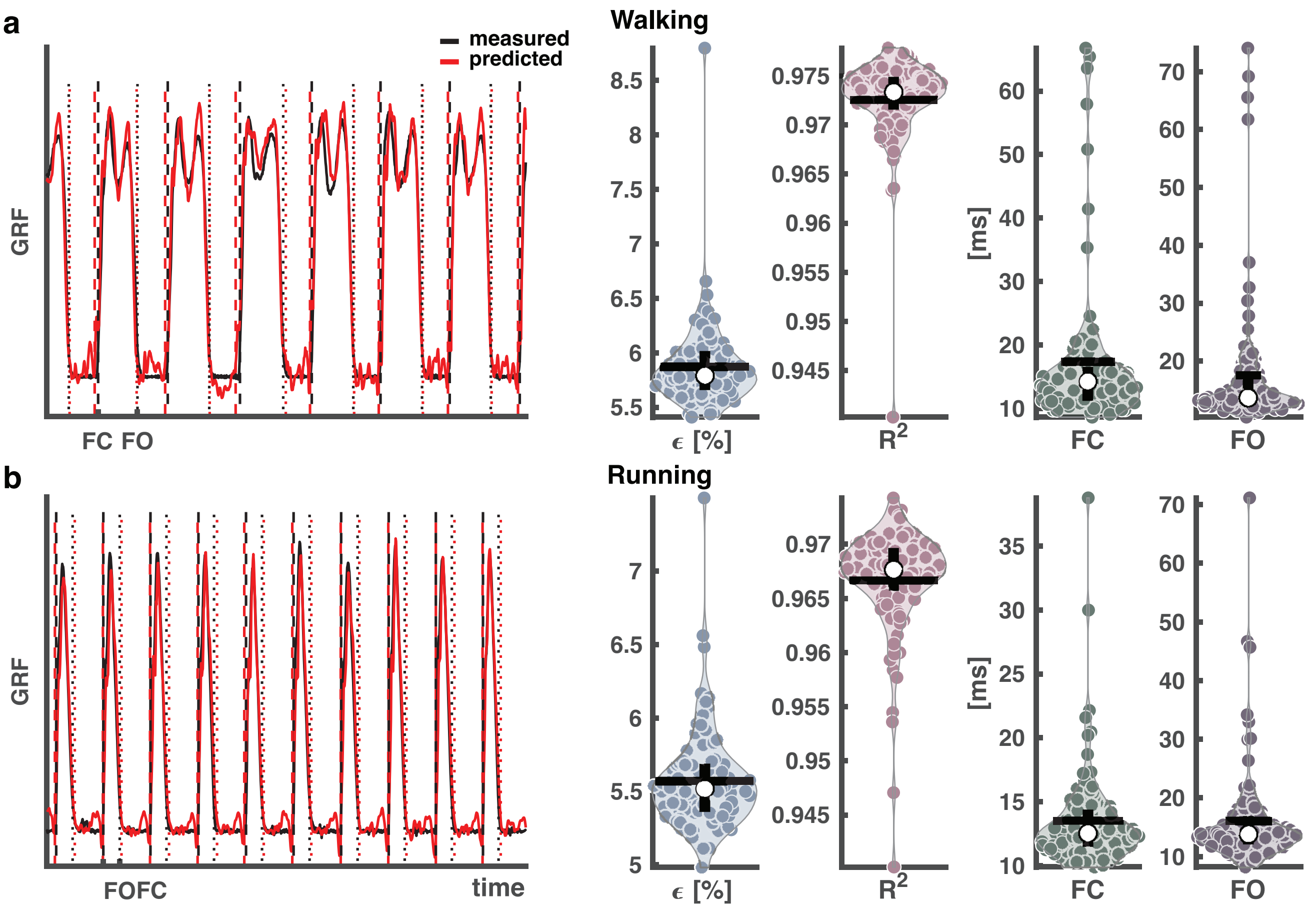


**Figure S1:** Output of the reservoir computer trained on segmented data and validated and tested on continuous data. a: Walking condition, b: Running condition. Left panels: Vertical ground reaction force (GRF) waveforms of four randomly selected strides from a random trial and a random selected

iteration out of the 100, with the measured vertical GRF waveforms in black and predicted in red. The vertical dashed lines represent the foot contact events (black: measured, red: predicted), and the vertical dotted one the foot off events (black: measured, red: predicted). Right side panels: Normalised root-mean squared error ($\epsilon$), coefficient of determination ($R^2$), mean absolute error of foot contact and foot off. The white dots in the violin-plots illustrate the medians. Black horizontal lines represent the mean and vertical black lines the 1st and 3rd quartiles. Every dot represents one of the 100 iterations. Abbreviations: GRF: ground reaction forces, FC: foot contact, FO: foot off.

As a final test of the validity of the machine learner, we ran the leave-M-out cross-validation for M = [1,2…6]. The leave-M-out iterations were all repeated 42 times to ensure no bias in the random selection of trials. The results can be found in Figure S2. The poor results found in the leave-one-out cross-validation stems from one trial with particularly large errors, here indicated as outliers.

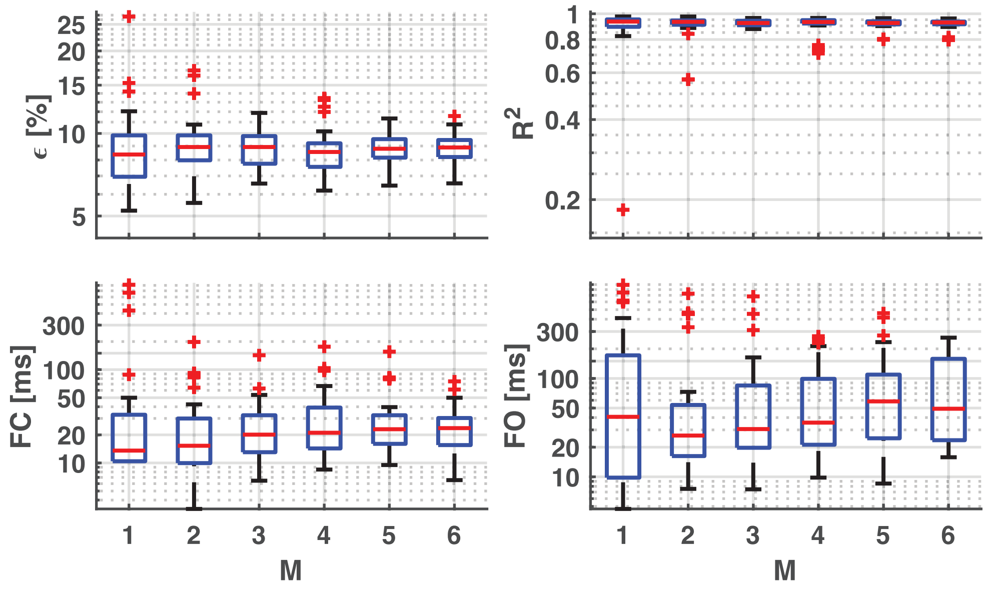


**Figure S2:** Result of the leave-M-out cross validation. A leave-M-out cross-validation with M ranging from 1 to 10 was carried out. Upper left panel: normalised root-mean squared error ($\epsilon$), upper right panel: coefficient of determination ($R^2$), lower left panel: mean absolute error of foot contact, and lower right panel: mean absolute error of foot off. Red horizontal lines indicate the median, the boxes represent the 25th and 75th percentile. Outliers are indicated with red plus. Note the large outliers in the M=1.
